# The effects of psychiatric history and age on self-regulation of the default mode network

**DOI:** 10.1101/342220

**Authors:** Stavros Skouras, Frank Scharnowski

**Author notes:** **Corresponding author:** Dr. Stavros Skouras. Postal address: Barcelonaβeta Brain Research Center, Carrer de Wellington 30, 08005 Barcelona.

## Abstract

Real-time neurofeedback enables human subjects to learn to regulate their brain activity, effecting behavioral changes and improvements of psychiatric symptomatology. Neurofeedback up-regulation and down-regulation have been assumed to share common neural correlates. Neuropsychiatric pathology and aging incur suboptimal functioning of the default mode network. Despite the exponential increase in real-time neuroimaging studies, the effects of aging, pathology and the direction of regulation on neurofeedback performance remain largely unknown. Using open-access analyses and real-time fMRI data shared through the Rockland Sample Real-Time Neurofeedback project (N=136), we first modeled neurofeedback performance and learning in a group of subjects with psychiatric history (n_a_=74) and a healthy control group (n_b_=62). Subsequently, we examined the relationship between up-regulation and down-regulation learning, the relationship between age and neurofeedback performance in each group and differences in neurofeedback performance between the two groups. Results show that in an initial session of default mode network neurofeedback with real-time fMRI, up-regulation and down-regulation learning scores are negatively correlated. Moreover, age correlates negatively with default mode network neurofeedback performance, only in absence of psychiatric history. Finally, adults with psychiatric history outperform healthy controls in default mode network up-regulation. Interestingly, the performance difference is related to no up-regulation learning in controls.

## Introduction

Neuroimaging research is contributing considerably to progress towards the apprehension of neurological and psychiatric disorders, by illustrating and characterizing their neural substrates. Current translational efforts are focused on validating non-invasive, neuroimaging-based diagnostic and therapeutic clinical applications. Out of the most innovative technological developments of our time, neurofeedback (NF) based on real-time functional magnetic resonance imaging (rt-fMRI) enables human subjects to learn to self-regulate their brain function, effecting behavioral changes and improvements of clinical symptomatology (Sitaram et al., 2017). Numerous recent studies have demonstrated therapeutic effects of rt-fMRI NF training on chronic pain (deCharms et al., 2005), addiction (Li et al., 2013; Hartwell et al., 2013), Parkinson’s disease (Subramanian et al., 2011), stroke (Robineau et al., 2017; Liew et al., 2016), tinnitus (Haller et al., 2010) and depression (Linden et al., 2012; Young et al., 2014; Young et al., 2017); consolidating the view that patients can learn to normalize abnormal patterns of brain activity that are associated with pathology (Sitaram et al., 2017; Stoeckel et al., 2014). Nevertheless, knowledge on the precise mechanisms underpinning the self-regulation of brain function is only nascent at present, and no conclusive theoretical framework for NF learning has been established (Sitaram et al., 2017). Especially in the context of clinical NF applications, the field is still lacking fundamental empirical evidence.

First, it has not been ascertained whether individuals with a history of psychiatric pathology are generally expected to perform equally to healthy participants in self-regulating brain function. Second, the way in which psychiatric pathologies affect specific cognitive requirements of a regulation task (e.g. mind wandering vs. focusing attention) is unknown. Third, key performance constraints, such as the effect of age, in relation to self-regulation of brain function and pathology, have not been investigated.

Here we address these three open issues regarding self-regulation of brain function, using the largest publicly available rt-fMRI NF repository, comprising data from healthy participants and psychiatric patients, during bidirectional self-regulation of the default mode network (DMN). The DMN is a large-scale cerebral network (Raichle et al., 2001) associated with a variety of brain functions, including perception (Kelly et al., 2008), attention (Weissman et al., 2006) and working memory (Mayer et al. 2010), which has become highly relevant for clinical applications (Zhang and Raichle, 2010; Brakowski et al., 2017; Mulders et al., 2015; Hamilton et al., 2015). Similarly to psychiatric pathology, aging also incurs suboptimal functioning of the human brain’s DMN (Whitfield-Gabrieli and Ford, 2012; Damoiseaux et al., 2008). Recent studies have demonstrated that DMN activity can be modulated through NF training (Harmelech et al., 2013; Megumi et al., 2015; Van De Ville et al., 2012). Thus, the DMN provides an optimal neural substratum for investigating the generic self-regulation of brain function.

Based on previous work assuming common neural mechanisms for up-regulation and down-regulation (Emmert et al., 2016) and proposing a generic neurofeedback learning network (Sitaram et al., 2017), we hypothesized that up-regulation and down-regulation learning scores would be positively correlated. Due to previously noted aberrant DMN function in pathological populations (Whitfield-Gabrieli and Ford, 2012), we expected that healthy participants would perform better than psychiatric patients in DMN self-regulation. Because of the impeding effects of aging on DMN function (Damoiseaux et al., 2008), we predicted that age would show a negative correlation with DMN self-regulation performance.

## Materials and Methods

### Participants

Young adults, aged 20 to 45 years (*M* = 30.94 years, *SD* = 7.32, *N* = 140; 58% female), took part in the study. Participants were residents of the Rockland County (New York, U.S.A.) and participated voluntarily in the Rockland Sample Real-Time Neurofeedback project, a large study aiming to create a sample with extensive neuropsychological and medical profiling, including several functional neuroimaging tasks (McDonald et al., 2017; Nooner et al., 2012). In order to include participants with a range of clinical and sub-clinical symptoms, minimally restrictive exclusion criteria were applied to exclude individuals with severe illnesses that would compromise compliance with experimental instructions (e.g. history of neoplasia requiring intrathecal chemotherapy or focal cranial irradiation, Global Assessment of Function < 50, history of psychiatric hospitalization, or suicide attempts requiring medical intervention). Modal psychiatric diagnoses were substance abuse and major depressive disorder. All subjects gave written informed consent; Institutional Review Board Approval was obtained for this project at the Nathan Kline Institute and at Montclair State University (Nooner et al., 2012). Subjects that presented a past or ongoing psychiatric diagnosis comprised the pathological group (*M* = 30.70 years; *SD* = 7.17; *n_a_* = 74; 55.4% female; 89.2% dextrous); subjects that had not presented any psychiatric diagnosis comprised the control group (*M* = 30.71 years; *SD* = 7.48; *n_b_* = 62; 50% female; 90.3% dextrous); and four unclassified subjects were excluded from further analysis.

### Experimental design

Participants completed a variety of assessments and functional neuroimaging tasks comprising the Enhanced Nathan Kline Institute-Rockland Sample (NKI-RS) protocol, described in detail elsewhere (McDonald et al., 2017; Nooner et al., 2012). The protocol included a six-minute resting state fMRI acquisition, followed by a twelve-minute neurofeedback task. Rt-fMRI data were used to derive neurofeedback performance and neurofeedback learning scores for each participant and for each direction of regulation, via general linear modeling.

Correlation testing was utilized to investigate the relation between DMN up-regulation neurofeedback learning and DMN down-regulation learning scores in the entire sample. Permutation testing was utilized to investigate differences in neurofeedback performance between the experimental group and the control group, for each direction of regulation. Nonparametric correlation testing was used to investigate the relation between neurofeedback performance and age, in each group.

To control for *Type I* error due to performing five individual statistical tests using data from the same sample, we used the Bonferonni correction method (Abdi, 2007) to adjust the typical significance level, *α* = 0.05 (see Statistical analysis). For the typical *Type II* error rate (*β* = 0.2), a priori computations of required sample size, using statistical power estimation software (G*Power v 3.1; Heinrich-Heine-Universität Düsseldorf RRID:SCR_013726; Faul et al., 2007; Faul et al., 2009), confirmed sufficient two-tail statistical power for effects of medium or large size using the entire sample (*d* > 0.487) and for effects of moderate or large size using only the pathological group (*d* > 0.660) or only the control group (*d* > 0.727).

### Resting state and neurofeedback task

A resting state acquisition preceded the neurofeedback task because it was required for the delineation of each participant’s DMN. During resting state, participants fixated on a white plus (+) sign centered on a black background for six minutes. During the neurofeedback task, stimuli comprised of the neurofeedback display illustrated in Figure 1B, which was implemented using a programming library (Vision Egg RRID:SCR_014589; Straw 2008) and is publicly available online from the ‘OpenCogLab Repository’ (GitHub; RRID:SCR_002630; https://git.io/vptev). All subjects participated in 12 counterbalanced, alternating trials of DMN up-regulation and down-regulation, of varying length (30 s, 60 s, 90 s). Each type of trial (e.g. up-regulation for 30 s) was repeated twice, once in the first and once in the second block of the session. Participants were instructed at the beginning of each trial to attempt to either let their mind wander (up-regulation), or to focus their attention (down-regulation), while attending to one out of four counterbalanced NF displays (Figure 1A). Counterbalancing was performed with regards to the following: 1) each trial featured a duration of either 30s, 60s or 90s; 2) each trial featured a central instruction, which was either “Focus” or “Wander”; 3) each trial featured the words “Focused” on the left and the word “Wandering” on the right or vice versa.

**Figure 1.**
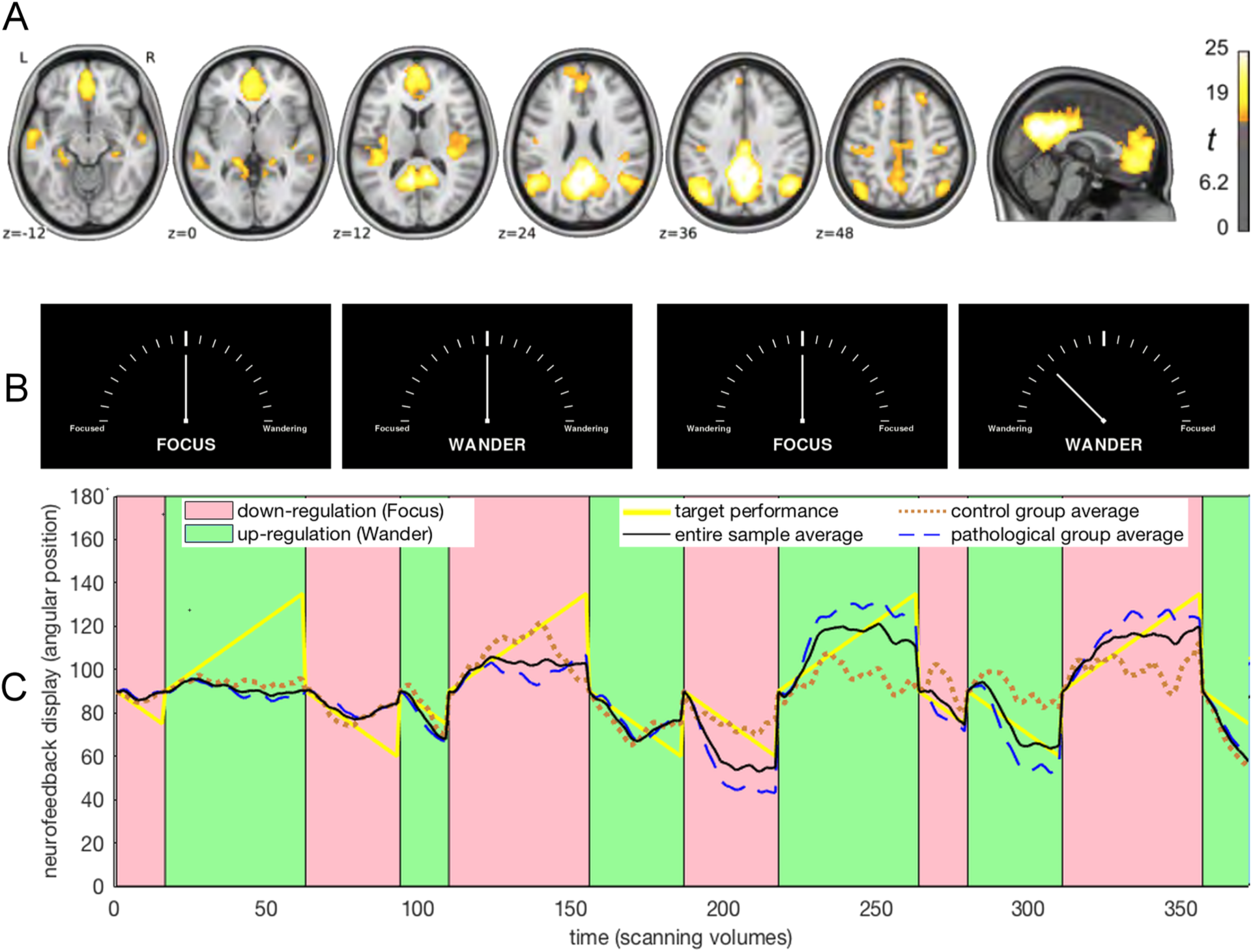
Modeling regulation performance. **A**, Spatial map of the average default mode network in the utilized repository (reprinted with permission, from McDonald et al. 2017). **B**, Each experimental trial featured one of the four counterbalanced displays resembling tachometers. At the beginning of each trial, the angular position of the needle pointer was set to 90° and was updated with every new functional volume according to real-time regulation performance; e.g. the rightmost display shows an angular position of 135°. **C**, The graph shows the average default mode network neurofeedback regulation performance for the pathological group (dashed blue line; N=74), for the control group (dotted brown line; N=62) and for the entire sample (black line; N=136), against target performance (yellow line), in one of four counterbalanced trial orders. It is apparent that on average, learning occurred for both up-regulation and down-regulation and that performance improved noticeably in the second half of the real-time fMRI scanning session.

### MRI data acquisition

Scanning was performed with a 3 T Siemens Magnetom TIM Trio scanner (Siemens Medical Solutions, Malvern, PA, USA) using a 12-channel head coil. Before the functional MR measurements, a fast localizer and high-resolution anatomical sequence were acquired. Anatomical images were acquired using a 3D T1-weighted MPRAGE (Mugler and Brookeman, 1990) GRAPPA (Griswold et al., 2002) sequence with an acceleration factor of 2 and 32 reference lines. 192 sagittal partitions were acquired, each of a 256×256 field of view (FOV), using a 2600 ms repetition time (TR), a 3.02 ms echo time (TE), 900 ms inversion time (TI) and 8° flip angle (FA), resulting in a 1 mm^3^ isomorphic voxel resolution. Functional images were acquired using echo planar imaging (EPI) with a TE of 30 ms, FA 90° and a TR of 2000 ms. Slice-acquisition was interleaved within the TR interval. The matrix acquired was 64×64 voxels with a FOV of 220 mm, resulting in an in-plane resolution of 3.4×3.4 mm^2^. Slice thickness was 3.6 mm with an interslice gap of 0.36 mm (30 slices). Images were exported over a network interface (Cox et al., 1995; LaConte et al., 2007). Physiological measures including galvanic skin response, pulse oximetry, rate of breathing and depth of breathing were measured during the functional acquisitions. Functional scanning included a six-minute resting state, prior to NF and other functional tasks using counterbalanced orders (McDonald et al., 2017).

### MRI data processing

Prior to the NF session, the DMN of each participant was delineated, based on the data from the resting state scan, using data preprocessing and support vector machines described elsewhere (McDonald et al., 2017). All neuroimaging data were subjected to the ‘Preprocessed Connectomes Project Quality Assessment Protocol’ (GitHub; RRID:SCR_002630; https://git.io/vptfA; Shehzad et al., 2015). A DMN template (Smith et al., 2009) was used to compute normalization transforms between MNI and anatomical space; white matter and cerebrospinal fluid signals were regressed out of the functional data and a support vector regression model was used to extract the DMN activation of each participant, for each TR, using software routines from AFNI (Analysis of Functional NeuroImages v 18.1.05; National Institute of Mental Health RRID:SCR_005927; Cox, 1996; LaConte et al., 2005) and FSL (FMRIB Software Library v 5.0; Oxford Centre for Functional MRI of the Brain RRID:SCR_002823; Zhang et al., 2001; Jenkinson et al., 2002; Jenkinson and Smith, 2001; Greve and Fischl, 2009; Greve and Fischl, 2009).

Data had been curated on and were accessed via the Collaborative Informatics and Neuroimaging Suite Data Exchange (COINS; Mind Research Network RRID:SCR_000805; Scott et al., 2011; Wood et al., 2014). The data featured a neurofeedback logfile for each participant, summarizing the measures of moment-to-moment neurofeedback performance derived by the real-time processing pipeline that was applied during data acquisition. The neurofeedback logfile for each participant was programmatically retrieved and read into computer memory using software (Matlab v 9.3; Mathworks RRID:SCR_001622). Structured variables were constructed to store all relevant information, regarding each participant’s and each trial’s characteristics and to perform hypothesis testing as illustrated in complete detail in the publicly available computer program developed for the analysis.

### Code accessibility

All original code developed for the analysis has been made available to any researcher for purposes of reproducing or extending the analysis, via the ‘NKI_RS_analysis’ public repository (GitHub; RRID:SCR_002630; https://git.io/vxddI). Following download, the analysis and progressive output can be viewed on any browser (HTML version) or replicated and modified interactively using the Live Editor in Matlab v 9.3 or higher (MLX version).

### Data accessibility

Neuroimaging data, further described in (McDonald et al., 2017), are available online via the Neuroimaging Informatics Tools Resources Clearinghouse (NITRC; RRID:SCR_003430). Diagnostic assessment data are available following the completion of a data transfer agreement with the Nathan Kline Institute (Nathan S. Kline Institute for Psychiatric Research; New York; USA, RRID:SCR_004334) via the Collaborative Informatics and Neuroimaging Suite Data Exchange (COINS; Mind Research Network RRID:SCR_000805).

### Statistical analysis

Standard general linear modeling was employed to quantify NF performance and learning. Using Pearson’s r as a metric of similarity, actual NF moment-to-moment regulation, as captured by the angular position of a needle on a tachometer-like NF display (Figure 1B), was compared to a linear vector representing the target performance of the experimental design (Figure 1C), producing a measure of NF performance for each of the 12 experimental trials. For each participant, average NF scores were computed separately for DMN up-regulation trials (wander), and DMN down-regulation trials (focus), in each of the two experimental blocks. NF learning score was computed as the improvement in NF regulation from the first to the second block of the session. NF performance score was computed as the average correlation between target performance and actual NF regulation in the second block of the session (following the introductory block of the session which comprised the participants’ very first practice experience of NF and of each trial type). Lilliefors test (Lilliefors, 1969) was used to assess normality and Grubbs’ test (Grubbs, 1969) was used for outlier detection. Parametric correlation testing was used to assess the relation between DMN up-regulation learning and DMN down-regulation learning score, because the variables were normally distributed. Non-parametric correlation testing was used to assess the relation between neurofeedback performance and age, because the distribution of age deviated from normality. Permutation testing was used to assess differences in neurofeedback performance scores between the experimental and control groups. Detailed documentation of the analysis, along with explanatory comments and an interactive program have been made available to facilitate replication and in support of the Open Science initiative. Five statistical tests were performed in total. To control for *Type I* error due to performing multiple statistical tests using data from the same sample, we used the Bonferonni correction method (Abdi, 2007) and set the adjusted two-tail significance level to *α* = 0.005.

## Results

### Direction of regulation

Lilliefors testing (Lilliefors, 1969) showed that NF up-regulation learning and down-regulation learning scores did not deviate from normality. Further quality assurance using Grubbs’ test for outlier detection (Grubbs, 1969) showed that no outliers were present in any utilized variables. Parametric correlation testing revealed that DMN NF up-regulation learning (*M* = 0.135, *SD* = 0.554, *N* = 136) and down-regulation learning scores (*M* = 0.189, *SD* = 0.587) were significantly negatively correlated, with a weak association explaining approximately 7% of the variance, *r(136)*=-0.258, *R^2^*=0.067, *P*=0.0024; Figure 2A.

**Figure 2.**
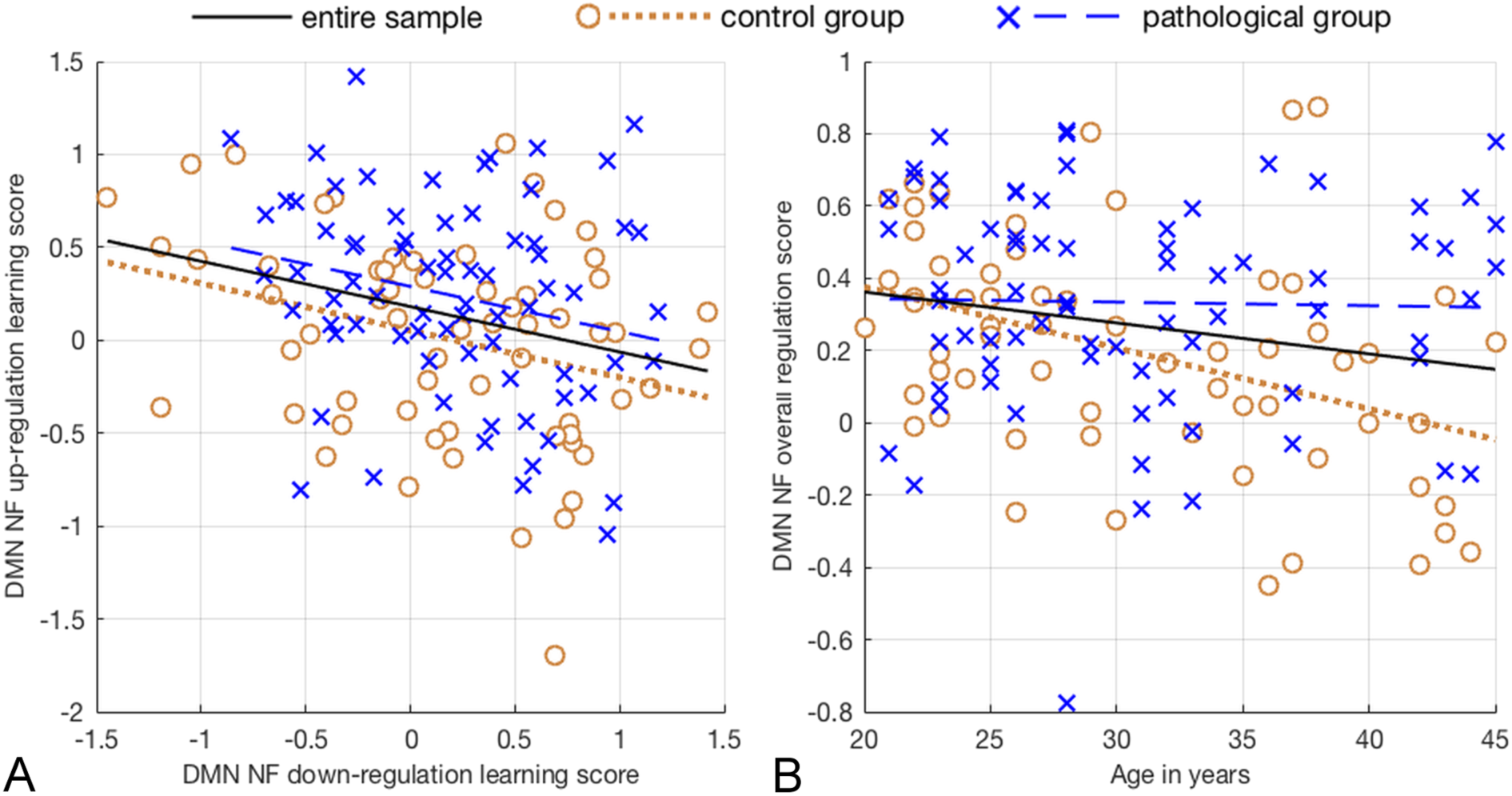
Illustration of the correlation tests performed. **A**, The figure shows lines of best fit for the pathological group (dashed blue line; N=74), the control group (dotted brown line; N=62) and the entire sample (black line; N=136). Default mode network neurofeedback up-regulation and down-regulation learning, are negatively correlated. **B**, Default mode network neurofeedback regulation performance decreases with age only in psychiatrically healthy adults.

### Age

Lilliefors testing showed that the distribution of age (*M* = 30.71 years, *SD* = 7.28, *N* = 136) deviated from normality. Non-parametric correlation testing using Spearman’s correlation coefficient revealed that in the pathological group, age (*M* = 30.70 years; *SD* = 7.17; *n_a_* = 74) had no effect on neurofeedback performance (*M* = 0.333, *SD* = 0.301), *r(74)* = −0.0711, *P* = 0.547. In the control group, age (*M* = 30.71 years; *SD* = 7.48; *n_b_* = 62) correlated negatively with DMN NF performance score (*M* = 0.195, *SD* = 0.312) with a moderate association that explained 17% of the variance, *r(62)* = −0.412, *R^2^* = 0.17, *P* = 0.0009; Figure 2B.

### Psychiatric pathology

Non-parametric independent samples t-tests, with 100,000 permutations per test, revealed that participants with a psychiatric history performed significantly better in DMN NF up-regulation (*M* = 0.383, *SD* = 0.420, *n_a_* = 74) than the control group (*M* = 0.159, *SD* = 0.467, *n_b_* = 62), *t*(134) = −2.951, *P* = 0.0036; *d* = 0.504; Figure 3A. Down-regulation performance was not significantly different between participants with a psychiatric history (*M* = 0.283, *SD* = 0.385) and the control group (*M* = 0.232, *SD* = 0.462), *t*(134) = −0.711, *P* = 0.479. The difference in up-regulation performance was due to a lack of learning, with regards to mind-wandering, in the control group; Figure 3B.

**Figure 3.**
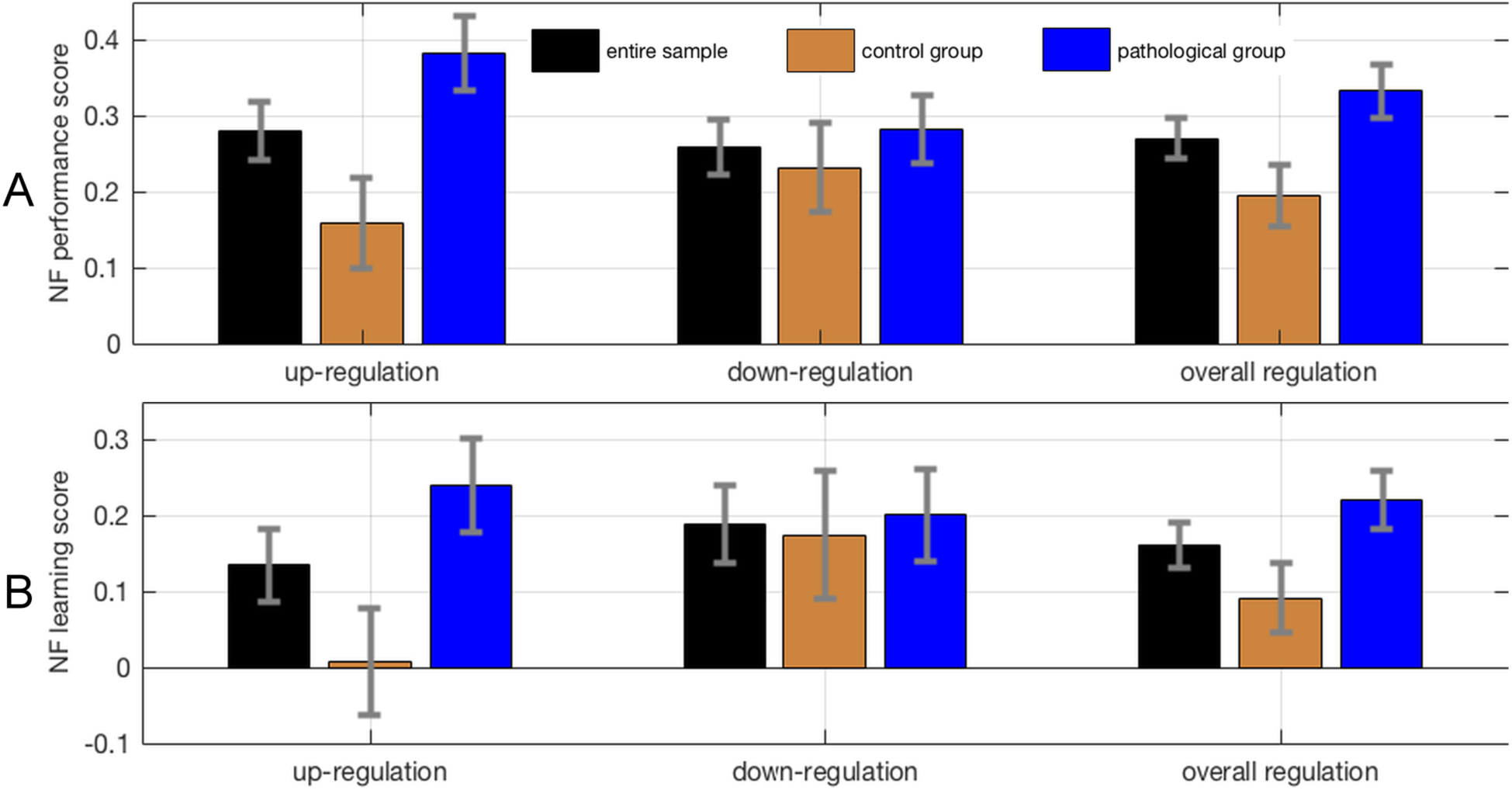
Neurofeedback scores across groups. **A**, Neurofeedback performance and neurofeedback learning scores for up-regulation, down-regulation and overall regulation. Scores are depicted separately for the control group, the pathological group and the entire sample. Error bars represent standard errors. Possible neurofeedback performance scores range from a value of −1 (corresponding to worst possible performance) to a value of 1 (corresponding to best possible performance). **B**, Possible neurofeedback learning scores range from a value of −2 (corresponding to worsening performance) to a value of 2 (corresponding to maximal learning).

## Discussion

In relation to existing questions of fundamental importance to self-regulating brain function, we have shown that initial learning of DMN self-regulation is influenced by age and psychiatric history. Specifically, during an initial, brief DMN rt-fMRI NF session, participants with a psychiatric history outperform healthy controls in DMN up-regulation. Further, age correlates negatively with the ability to self-regulate DMN function, only in participants without psychiatric history. Finally, up-regulation learning and down-regulation learning scores are negatively correlated in both patients and healthy controls.

A negative effect of age on DMN self-regulation is not surprising, given reduced resting-state brain activity (Kelly et al., 2008) and connectivity (Sambataro et al., 2010; Dennis and Thompson, 2014) within the DMN in normal aging. The DMN also comprises the primary locus of earliest amyloid deposition due to aging in cognitively normal individuals (Kelly et al., 2008; Palmqvist et al., 2017). Although it does not affect cognitive performance, such a neurobiological burden poses limitations on the neural efficiency of the DMN, leading to functional connectivity changes (Palmqvist et al., 2017) and age-related decline in task-related modulation of DMN activity (Sambataro et al., 2010). Our finding indicates that in healthy subjects, age explains 17% of the variation in NF self-regulation scores. This strong effect is even more remarkable considering that the age range in our study was limited to young adults (20-45 years of age), similarly to most NF studies to date. Hence, age must be considered and explicitly modeled when assessing the ability to self-regulate brain function.

Although no previous rt-fMRI study investigated age effects on rt-fMRI NF performance, our finding corroborates complementary evidence, regarding self-regulation, from electrophysiological biofeedback and neurofeedback studies. Heart rate variability biofeedback performance has been reported to correlate negatively with age and the effects of biofeedback self-regulation training on cardiovascular measures were less pronounced in older subjects (Lehrer et al., 2006). EEG theta amplitude up-regulation was also illustrated to be noticeably better in young compared to old subjects, although that study did not investigate performance differences between age groups via statistical testing (Wang and Hsieh, 2013).

DMN activity has been linked primarily to occipital alpha band activity (Jann et al., 2010) that is known to decrease with age (Breslau et al., 1989), substantiating a plausible physiological basis for the effects observed here. It should be noted that despite the evidence for decreased self-regulation performance in older subjects, the beneficial effects of neurofeedback and biofeedback self-regulation training on cognitive and respiratory function, are stronger in elders (Lehrer et al., 2006; Wang and Hsieh, 2013), suggesting the suitability of self-regulation training as a promising intervention to support healthy aging.

A surprising finding from our study relates to the absence of an age effect in the pathological group. In cohort with the complementary finding of better performance in the pathological group, our results suggest that good self-regulation performance is not necessarily reflecting healthy brain function. Higher up-regulation performance in the pathological group is supported by pathologically altered DMN connectivity at baseline (Broyd et al., 2009) and may be associated to reduced DMN coherence, pointing to higher functional stability of the DMN in the control group and in healthy aging. These observations regarding initial DMN regulation performance suggest that generic hypotheses of reduced performance in pathological populations need to be revisited.

It is worth considering that the capacity to show improvement in NF performance is contingent upon aberrant baseline activity. That is, when NF is used to normalize brain activity in patients that present moderately abnormal, aberrant activity, the patients will probably perform better than controls that present normal, stable activity patterns. On the other hand, in the face of severe pathology that impairs fundamental learning capacities, the ability to improve NF performance will be impaired as well, although this was not the case for the high-functioning pathological group of the present study.

The third important finding of our study is that initial learning of DMN self-regulation is unidirectional, with the amount of learning in one direction of regulation opposing learning in the other direction of regulation and explaining 7% of its variation. In this study, up-regulation was associated with mind-wandering strategies and down-regulation with focusing attention. Thus, our findings suggest that learning to self-regulate depends on task-specific performance constraints rather than a more generic ability to self-regulate. This may partly explain why the search for general predictors of successful neurofeedback learning has not been successful so far and why the proportion of up to 30% of participants that fail to self-regulate effectively, even following extensive practice (Harmelech et al., 2015), varies between targeted brain regions and paradigms (Sitaram et al., 2017). In order to design improved experimental and clinical protocols, it is critical to consider the direction of regulation, task-specific peculiarities, and individual predispositions.

It remains possible that the present findings may be influenced by unaccounted factors that are not homogeneously distributed across the two experimental groups or across the age range of the sample. These may include experience with types of training that can exert a covert influence on the ability to self-regulate physiological processes; e.g. meditational practice, musical training, acting experience etc. (Gruzelier, 2014a, 2014b). It is possible that, on average, the pathological group had received more training in self-regulation of cognitive and physiological functions through participation in cognitive behavioral therapy programs. Moreover, it remains possible that our findings are limited only to the DMN and only to initial sessions of self-regulation. Future research should systematically investigate learning effects in cardinal brain networks (i.e. default mode, executive control and salience networks) across several sessions using identical protocols.

In conclusion, we have illustrated that participants with a psychiatric history can perform better than healthy controls in self-regulating brain function. The ability to up-regulate brain function appears to decrease with age only in the absence of psychiatric history, and good performance in learning to up-regulate the DMN through mind-wandering does not imply equally good performance in down-regulating the DMN through focusing attention. Although it remains to be determined to what extent these findings are specific to the DMN and to initial sessions of self-regulation through rt-fMRI NF, our findings contribute to the furtherment of neurophysiological knowledge and our understanding of the neural mechanisms underpinning self-regulation of brain function specifically. We anticipate that our findings will solidify the foundations for more systematic modeling of the self-regulation of brain function and assist in the design of advanced technologies and experimental protocols for clinical applications.

## Acknowledgements

This work was supported by the European Union’s Horizon 2020 research and innovation programme under the Marie Sklodowska-Curie action grant agreement No 707730, the Foundation for Research in Science and the Humanities at the University of Zurich (STWF-17-012), the Baugarten Stiftung, and the Swiss National Science Foundation (BSSG10_155915, 32003B_166566). Principal support for the Rockland Sample Real -Time Neurofeedback project was provided by the NIMH BRAINS R01-MH101555 (PI Craddock). The authors thank Cameron Craddock for insightful discussions. The authors declare no competing financial interests.

